# Evaluation of *trans*- and *cis*-4-[^18^F]fluorogabapentin for brain PET imaging

**DOI:** 10.1101/2023.09.01.555353

**Authors:** Yu-Peng Zhou, Marc D. Normandin, Vasily Belov, Marina T. Macdonald-Soccorso, Sung-Hyun Moon, Yang Sun, Georges El Fakhri, Nicolas J. Guehl, Pedro Brugarolas

## Abstract

Gabapentin, a selective ligand for the α2δ subunit of voltage-dependent calcium channels, is an anticonvulsant medication used in the treatment of neuropathic pain, epilepsy and other neurological conditions. We recently described two radiofluorinated derivatives of gabapentin (*trans-*4-[^18^F]fluorogabapentin, [^18^F]tGBP4F, and *cis-*4-[^18^F]fluorogabapentin, [^18^F]cGBP4F) and showed that these compounds accumulate in the injured nerves in a rodent model of neuropathic pain. Given the use of gabapentin in brain diseases, here we investigate whether these radiofluorinated derivatives of gabapentin can be used for imaging α2δ receptors in the brain. Specifically, we developed automated radiosynthesis methods for [^18^F]tGBP4F and [^18^F]cGBP4F and conducted dynamic PET imaging in adult rhesus macaques with and without preadministration of pharmacological doses of gabapentin. Both radiotracers showed very high metabolic stability, negligible plasma protein binding and slow accumulation in the brain. [^18^F]tGBP4F, the isomer with higher binding affinity, showed low brain uptake and could not be displaced whereas [^18^F]cGBP4F showed moderate brain uptake and could be partially displaced. Kinetic modeling of brain regional time-activity curves using a metabolite-corrected arterial input function shows that a 1-tissue compartment model accurately fits the data. Graphical analysis using Logan or multilinear analysis 1 produced similar results as compartmental modeling indicating robust quantification. This study advances our understanding of how gabapentinoids work and provides an important advancement towards imaging α2δ receptors in the brain.

## INTRODUCTION

Gabapentin (GBP, **Fig. 1)** targets the α2δ subunit of voltage-gated calcium channels (VGCC) expressed throughout the central and peripheral nervous systems (CNS and PNS)^1-4^. Four subunits of α2δ (α2δ-1 through 4) have been identified and their expression is altered in multiple diseases^5^. For instance, the expression of α2δ-1 in the PNS or CNS is greatly increased in models of neuropathic pain^6-12^, neuroinflammation^13,14^ and addiction^15-20^ whereas α2δ-1 protein is downregulated in an animal model of epilepsy^21^. In addition, mutations in α2δ-2 are associated with seizures in animals and humans^22,23^. As such, pharmacological modulation of α2δ with gabapentin results in clinical improvement of conditions such as partial seizures and neuropathic pain^24,25^.

**Figure 1.**
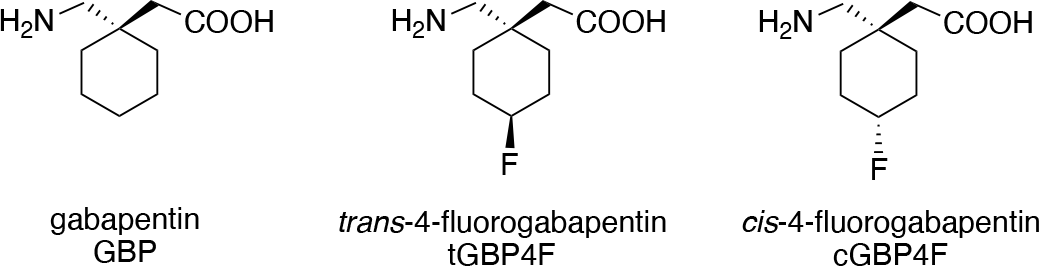
Structures of gabapentin (GBP, 2-(1-(aminomethyl)cyclohexyl)acetic acid) and fluorinated derivatives *trans*-4-fluorogabapentin (tGBP4F, 2-((1*R*,4*R*)-1-(aminomethyl)-4-fluorocyclohexyl)acetic acid) and *cis*-4-fluorogabapentin (cGBP4F, 2-((1*S*,4*S*)-1-(aminomethyl)-4-fluorocyclohexyl)acetic acid). tGBP4F indicates that the fluorine is in *trans*- to the -CH_2_NH_2_ and cGBP4F indicates that the fluorine is in *cis*- to the -CH_2_NH_2_.

Given the role of α2δ in multiple CNS and PNS diseases, we hypothesized that imaging α2δ with a radiolabeled version of gabapentin would be valuable for understanding these diseases and could potentially serve as a diagnostic tool. In our prior work, we developed two radiofluorinated derivatives of gabapentin (*cis*- and *trans*-[^18^F]4-fluorogabapentin or [^18^F]cGBP4F and [^18^F]tGBP4F, **Fig. 1**) and found that these compounds bind to α2δ-1 with very different affinities (the IC_50_ values measured in a competitive radioligand binding assay were 24 nM for [^18^F]tGBP4F and 2400 nM for [^18^F]cGBP4F)^26^. We also found that these compounds accumulate 1.6- to 2-fold higher in the injured nerves than uninjured nerves of a rat model of neuropathic pain, which is consistent with the higher protein expression in these tissues.

Given the important role of α2δ subunits in CNS diseases, we decided to examine the [^18^F]tGBP4F and [^18^F]cGBP4F for brain imaging. Since the affinity of [^18^F]cGBP4F for α2δ-1 is likely insufficient to achieve specific binding *in vivo*, imaging with this isomer may provide the opportunity to control for non-specific binding. The paradigm of using active and inactive isomers in PET to assess specific binding has been previously used^27^. Although these tracers have a low predicted partition coefficient (cLogP = -1.267), which suggests poor brain penetration, it is known that gabapentin enters the brain via the LAT-1 transporter^28^ and, thus, these tracers may have suitable brain penetration. To investigate the potential value of these tracers for brain PET imaging, we used non-human primates (NHPs) as their large brain size and similarities with the human brain make them excellent models to study brain tracers. Furthermore, to produce enough tracer dose for large animal studies, fully automated radiosynthesis methods were developed consistent with current good manufacturing practices (cGMP) and radiation safety regulations.

## RESULTS

### Automation of the radiosynthesis of [^18^F]tGBP4F and [^18^F]cGBP4F

The radiosynthesis of PET tracers [^18^F]tGBP4F and [^18^F]cGBP4F has been previously reported (**Fig. 2**)^26^. Here, automated radiosynthesis methods were developed using a GE TRACERlab FX2N two-reactor synthesis module. Before starting the radiosynthesis, the reagents were loaded into the different vials as shown in **Supplemental Figure S1**. In the first step, [^18^F]fluoride was trapped in a QMA cartridge, eluted with H_2_O/MeCN/kryptofix/K_2_CO_3_ into reactor 1 and dried under reduced pressure and heating. The isomerically pure precursor **1a** or **1b** dissolved in DMSO was added to reactor 1 and radiofluorination was carried out at 140 ºC for 20 min forming intermediate **2a** or **2b**, respectively (**Fig. 1**). Intermediate **2a** or **2b** was diluted in water and trapped on an HLB cartridge. Next, the intermediate **2a** or **2b** was eluted from the cartridge using acetonitrile and transferred into reactor 2 for hydrolysis with NaOH and deprotection with HCl. Since the final product is hydrophilic, [^18^F]tGBP4F or [^18^F]cGBP4F was isolated by semi-preparative HPLC purification by using a mobile phase compatible for injection containing 5% EtOH in phosphate buffer. A typical run starts from 37 GBq (1.0 Ci) of [^18^F]fluoride and yields 1.6 GBq (43 mCi) of [^18^F]tGBP4F (non-decay corrected radiochemical yield: 4.3 ± 2.2 %, n = 13) or 0.52 GBq (14 mCi)

**Figure 2.**
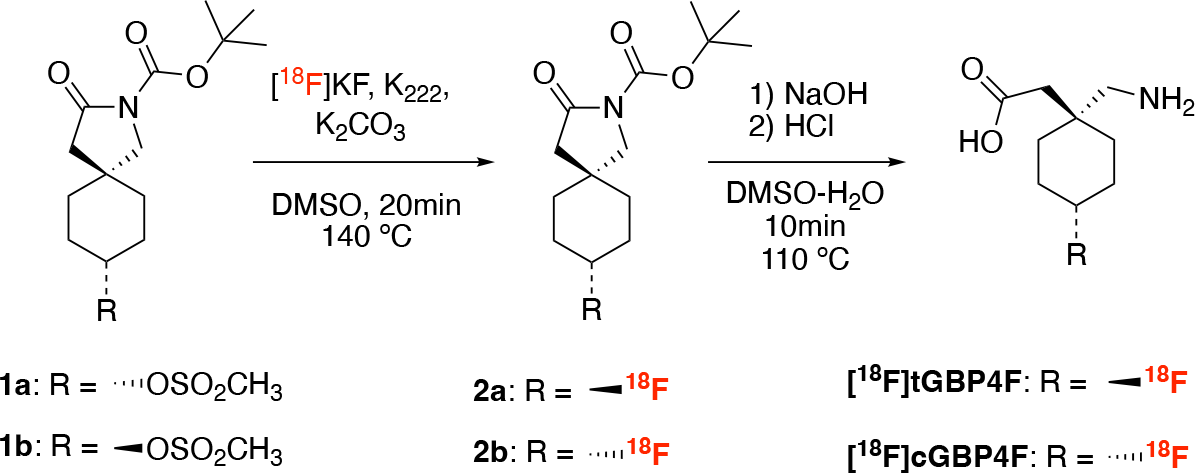
Automated production of [^18^F]tGBP4F and [^18^F]cGBP4F. **A**. Radiochemical synthesis of [^18^F]tGBP4F and [^18^F]cGBP4F.

[^18^F]cGBP4F (non-decay corrected radiochemical yield: 1.4 ± 1.2 %, n = 3) in ∼120 min. The radiochemical purity of [^18^F]tGBP4F or [^18^F]cGBP4F was calculated by integrating all the radioactive peaks on analytical radioHPLC and found to be >99% (**Sup. Fig S2**). Since these compounds have negligible UV absorption, the identification of the tracer (*cis*- or *trans*-) was not based on coinjection but on the identity of the precursor. The molar activities for[^18^F]tGBP4F and [^18^F]cGBP4F were estimated to be greater than 37 GBq/μmol (1 Ci/μmol) at end of synthesis.

### [^18^F]tGBP4F and [^18^F]cGBP4F have negligible plasma protein binding and are not metabolized

Two rhesus macaques underwent paired baseline and blocking PET scans with either [^18^F]tGBP4F or [^18^F]cGBP4F within 3 months of each other. For blocking scans, animals were injected with gabapentin (4.7 mg/kg for the [^18^F]tGBP4F scan; 5.0 mg/kg for the [^18^F]cGBP4F scan) 30 min before radiotracer injection.

Serial arterial blood samples were collected every 30 seconds upon radiotracer injection and decreased in frequency to every 15 minutes toward the end of the scan. Selected blood samples were analyzed for radioactivity concentrations in whole blood (WB) and plasma (PL). Selected plasma samples were analyzed by radio-HPLC to measure the parent fraction in plasma (PP). The plasma free fraction (*f*_*p*_) was also measured. The WB and parent in PL curves peaked at ∼3.5 min after radiotracer injection and the decline following the peak was best described by a biexponential curve (**Fig. 3** and **Sup. Fig. S3**). The half-lives of the rapid and slow components of the parent in plasma are shown in **Table 1** for each tracer ([^18^F]cGBP4F and [^18^F]tGBP4F) and condition (baseline and blocking). The clearance was 147 min for the *cis*- and 88 min for the *trans*- and it became faster under blocking conditions (99.7 min and 79.0 min, respectively).

**Figure 3.**
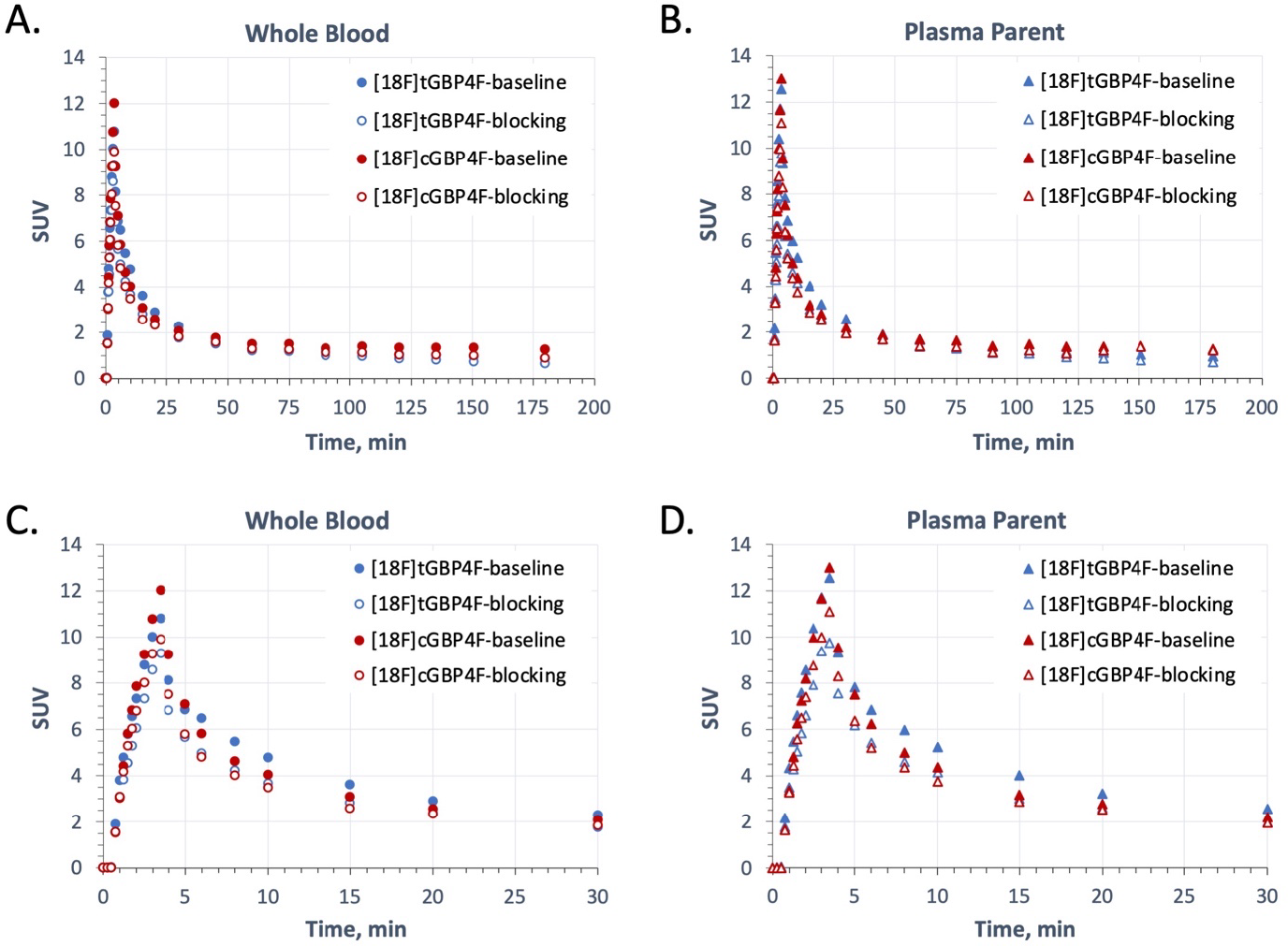
Blood and plasma concentrations of [^18^F]tGBP4F and [^18^F]cGBP4F. **A**. Time courses of radioactivity concentrations in arterial whole blood under baseline and blocking conditions. **B**. Time courses of parent in plasma under baseline and blocking conditions. **C**. and **D**. Magnification of the first 30 min of **A** and **B**, respectively.

**Table 1.** Plasma kinetic parameters from biexponential model fits.

| Tracer / condition | Fast component half-life (min) | Slow component half-life (min) | R<sup>2</sup> |
| --- | --- | --- | --- |
| cGBP4F / baseline | 2.86 | 147 | 0.989 |
| cGBP4F / blocking | 2.64 | 99.7 | 0.989 |
| tGBP4F / baseline | 4.54 | 88.5 | 0.994 |
| tGBP4F / blocking | 3.40 | 79.0 | 0.994 |

The plasma free fraction (*f*_p_) averaged 1.01 ± 0.03 across studies for each radiotracer indicating negligible plasma protein binding. The WB to PL ratios reached equilibrium within 1 min and remained stable at near 0.9 for all scans (**Sup. Fig. S4**). RadioHPLC analysis of plasma samples suggested that both radiotracers undergo minimal *in vivo* metabolism (%PP range from 95% to 98% at 180 min post injection (**Sup. Fig. S5**)).

### Brain PET imaging

Summed PET images of the monkey brains from 45-90 min show moderate brain uptake for [^18^F]cGBP4F and low brain uptake for [^18^F]tGBP4F (**Fig. 4**). Time-activity curves (TACs) were extracted for nineteen brain ROIs as described in the methods section. The regional TACs for [^18^F]cGBP4F baseline scan peaked around 3.8 min (whole brain SUV = 0.55), followed by a sharp decrease (SUV = 0.45 at 9 min) and a slow accumulation in the brain (SUV at 180 min = 0.7). Similarly, [^18^F]tGBP4F TACs showed an initial peak (SUV = 0.4 at 3.8 min), followed by a decrease (SUV = 0.25 at 13 min) and a very slow accumulation in the brain (SUV = 0.28 at 180 min). Within the brain, the areas with highest uptake were the amygdala, insula, nucleus accumbens and cortex for both tracers and the areas with lowest uptake were the thalamus, cerebellum and white matter.

**Figure 4.**
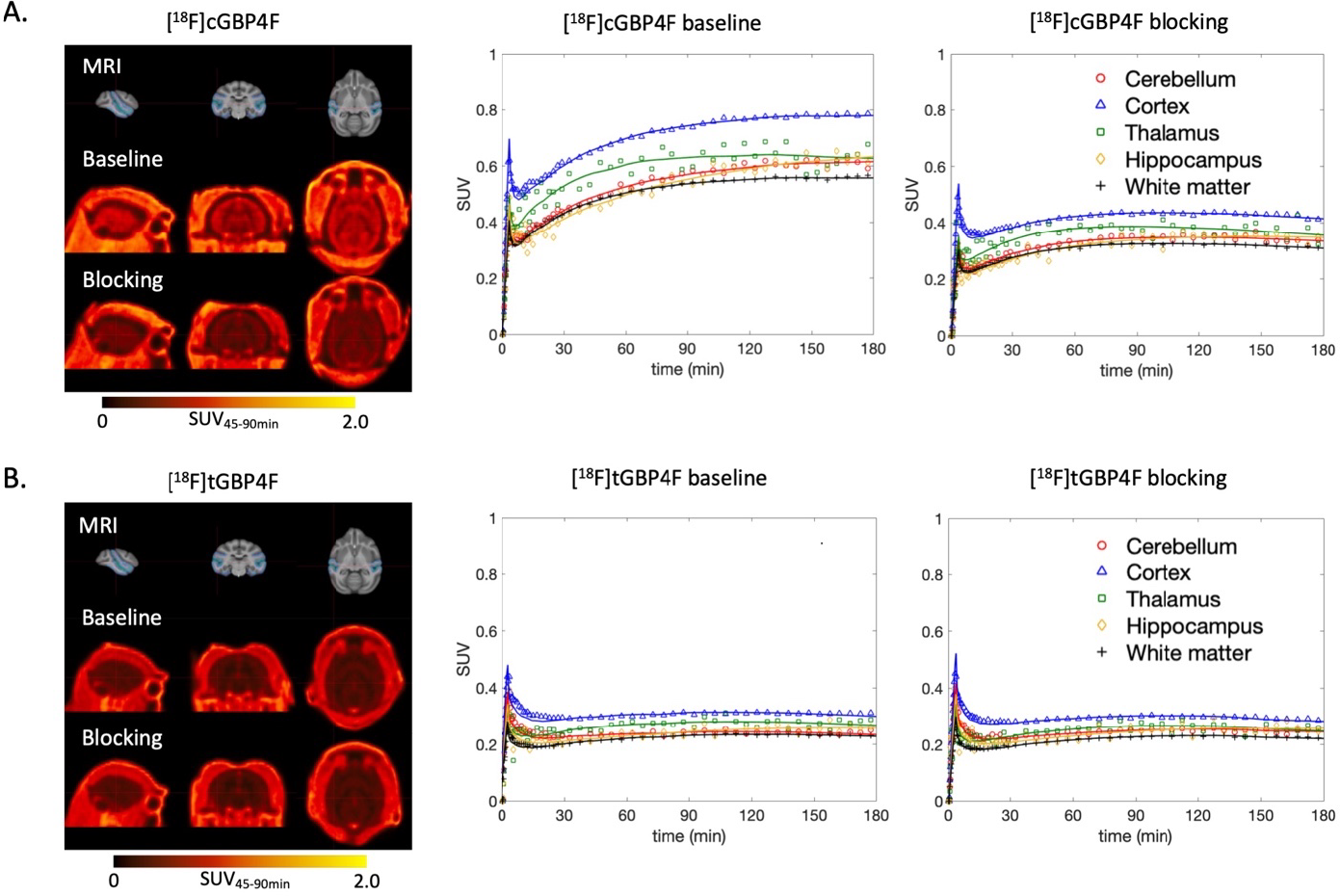
PET images and time activity curves of [^18^F]cGBP4F and [^18^F]tGBP4F under baseline and blocking conditions. **A**. PET SUV images showing the cerebral distribution of [^18^F]cGBP4F under baseline and blocking conditions and corresponding regional time activity curves. **B**. PET SUV images showing the cerebral distribution of [^18^F]tGBP4F under baseline and blocking conditions and corresponding regional time activity curves. Solid lines represent fitting from a 1TCM.

In the case of [^18^F]cGBP4F, preinjection of 5 mg/kg of non-radioactive gabapentin resulted in a visible reduction in brain SUV form 45-90 min which is also evidenced by a reduction in the TACs across all brain regions (**Fig. 4A**). For [^18^F]tGBP4F, pre-injection of 4.7 mg/kg of non-radioactive gabapentin did not result in an obvious change in tracer uptake or in the regional TACs (**Fig. 4B**). In order to better quantify the blocking effects compartmental modeling was performed.

### Modeling of the imaging data

Kinetic modeling of the regional TACs was performed using 1-tissue and 2-tissue compartment models (1TCM and 2TCM) with fixed and variable vascular contribution using the whole scan duration (180 min) and after truncating the data to 120 and 90 min. A 1TCM with vascular contribution as a model parameter accurately fit the full TACs for both tracers and both conditions (**Fig. 4**).

For [^18^F]cGBP4F under baseline conditions, *V*_T_ values ranged from 0.58 mL/cc for the amygdala to 0.34 mL/cc for the white matter (**Fig. 5**) and *K*_1_ values ranged from 0.0059 mL/min/cc for the occipital gyrus to 0.0029 mL/min/cc for the hippocampus. Under blocking conditions, the *V*_T_ and *K*_1_ values for the whole brain decreased -36% and -35%, respectively. The correlation between changes in *V*_T_ and changes in *K*_1_ across all brain regions (Pearson R = 0.479) suggests that the reduction in *V*_T_ may be driven by saturation of the LAT-1 transporter by cold gabapentin.

**Figure 5.**
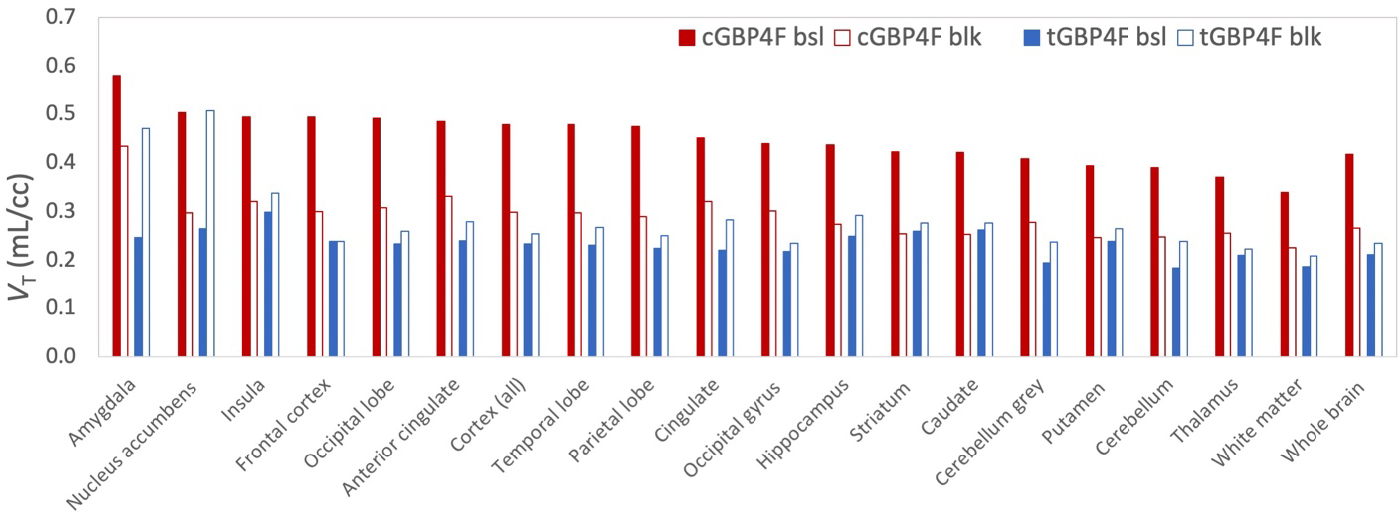
Regional *V*_*T*_ estimates obtained from the 1TCM for both tracers and conditions.

In the case of [^18^F]tGBP4F, *V*_T_ values ranged from 0.30 mL/cc for the insula to 0.18 mL/cc for the cerebellum (**Fig. 5**) and *K*_1_ values ranged from 0.0021 mL/min/cc for the occipital gyrus to 0.0010 mL/min/cc for the amygdala. Under blocking conditions, the *V*_T_ and *K*_1_ values for the whole brain increased by 11% and 16%, respectively. However, given the low *V*_T_ and *K*_1_ values, it is difficult to assess the significance.

Graphical analysis using the Logan or multilinear analysis 1 (MA1) resulted in good linearization after 60 min (t* = 60 min) (**Sup. Fig. S6**). *V*_T_ values calculated with MA1 (**Fig. 6**) were in better agreement with *V*_T_ obtained from the compartmental analysis compared to those calculated with Logan (**Sup. Fig. S7**). Logan *V*_T_ suffered from a small underestimation and larger standard deviation (**Table 2**). Truncating the data to 120 or 90 min resulted in less robust *V*_T_ estimates and greater variability, as it would be expected given the slow kinetics.

**Figure 6.**
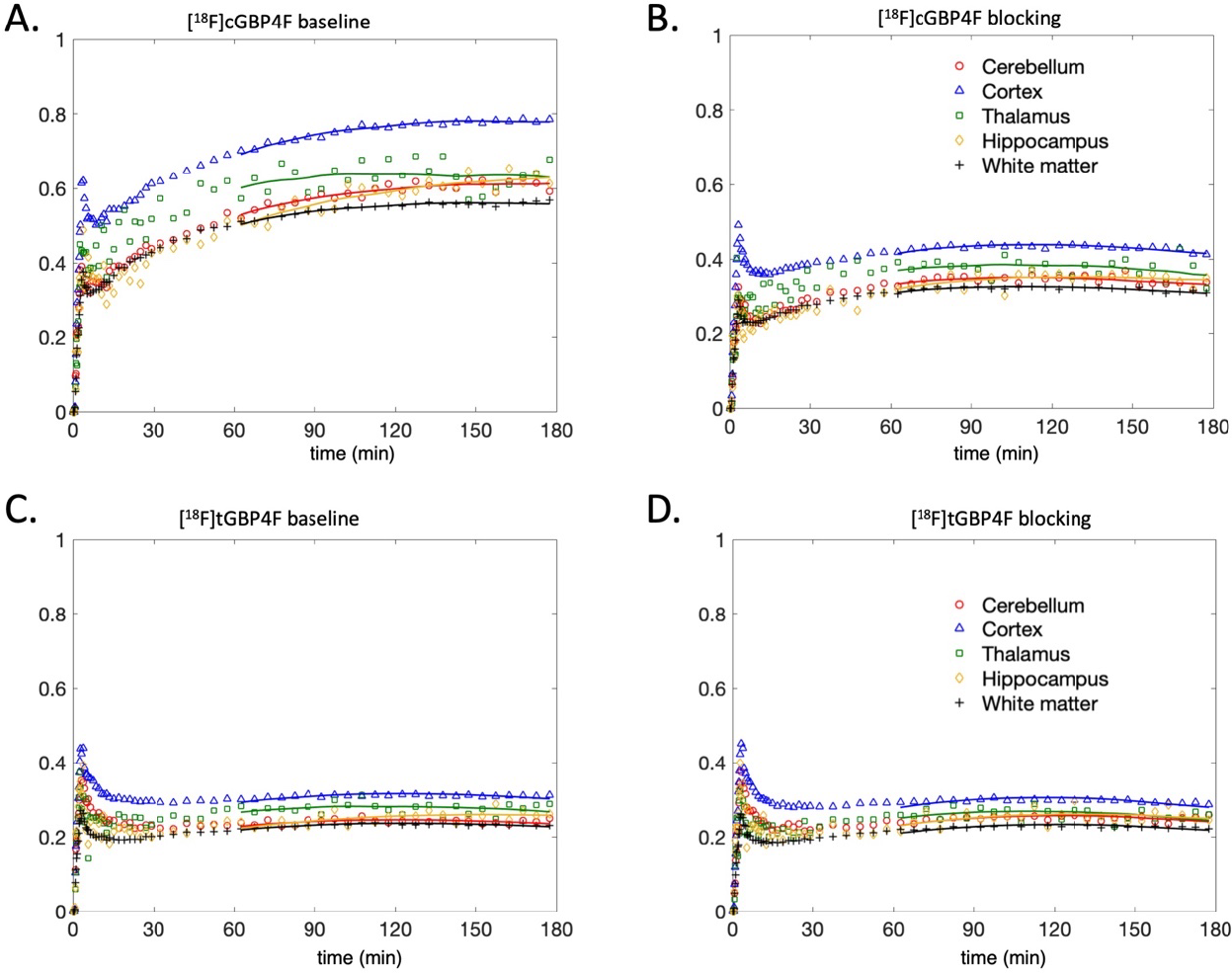
MA1 graphical analysis plots for [^18^F]cGBP4F and [^18^F]tGBP4F under baseline and blocking conditions. End time (t_end_) of 180 min and a t* of 60 min were used for MA1 fits. MA1 fits to the data over the t* to t_end_ time interval is depicted for each representative ROI.

**Table 2.** Percent difference *V*_T_ *vs. V*_T_ 1T2kv.

| Tracer | Logan (180 min) | MA1 (180 min) |
| --- | --- | --- |
| cGBP4F | $-0.5 \pm 7.9$ | $3.2 \pm 3.3$ |
| tGBP4F | $-4.2 \pm 13.1$ | $0.8 \pm 8.1$ |

## DISCUSSION

The present study focuses on the evaluation of [^18^F]tGBP4F and [^18^F]cGBP4F, which have previously been found to have high and moderate affinity, respectively, to α2δ subunits^26^. Since α2δ subunits play a crucial role in various brain diseases, the development of radioligands that can image these proteins in the brain is of great interest. In this study, an automated radiolabeling method was developed to produce sufficient quantities of the radioligands with high molar activity for NHP studies.

The NHP studies showed that the radioligands are highly stable in blood and free of plasma protein binding. Their brain uptake was found to be moderate to low, and their accumulation in the brain very slow. The tracer kinetics across brain regions could accurately be described using a 1-tissue compartmental model which produced *V*_T_ values in the range 0.2 to 0.6 mL/cc for both tracers. When preinjected with gabapentin, [^18^F]cGBP4F showed a -35% reduction in *V*_T_ and [^18^F]tGBP4F showed a 16% increase. The different effect upon drug challenge is unexpected may be due to different affinities to the LAT-1 transporter, partial blocking of the receptor or other reasons. In the absence of test-retest studies, we cannot rule out experimental variability, however, given that the isomer with the higher binding affinity ([^18^F]tGBP4F) had low brain uptake and could not be blocked and that the tracer with higher brain uptake and partially blockable ([^18^F]cGBP4F) had low binding affinity, it did not seem warranted to pursue additional imaging studies in nonhuman primates with these compounds. Nevertheless, this study provides some novel insights about how gabapentinoids work. For example, it is remarkable that the orientation of a fluorine atom in the 4-position can have, not only a large effect on the binding affinity to the α2δ protein as previously described^26^, but also result in markedly different brain uptake despite the two isomers having very similar physicochemical properties and similar blood input functions. Furthermore, these novel compounds may still find value as therapeutics given the slow kinetics.

In conclusion, while the investigation of [^18^F]tGBP4F and [^18^F]cGBP4F provides new information on the brain uptake and kinetics of GBP derivatives, the goal of imaging α2δ in the brain using PET remains elusive. Nevertheless, the findings of this study can be leveraged towards the development of novel tracers with improved imaging properties and potentially as therapeutic agents for brain diseases involving α2δ subunits.

## METHODS

### Materials

Commercially available reagents were purchased from Sigma-Aldrich, Acros, Alfa-Assar or TCI and used as received.

### Automated radiosynthesis of PET tracers

To start, target water from cyclotron was send to the QMA cartridge where [^18^F]fluoride was trapped and eluted to reactor 1 by K_222_/K_2_CO_3_ eluent. After drying the [^18^F]fluoride, the isomerically pure precursor (**1a** or **1b**) dissolved in DMSO was added to the reactor and then heated to 140°C for 20 min. Sterile water (2 x 6mL) was added to the reactor 1 and the reaction mixture was sent to an HLB cartridge (see **Fig 1**). The labeled intermediate (**2a** or **2b**) was trapped on the HLB cartridge and eluted into reactor 2 using 2 mL of acetonitrile. NaOH and HCl solutions were added to reactor 2 and heated to 120°C for 5 min, respectively. Additional NaOH was added to neutralize the solution and vacuum was applied to remove acetonitrile. 2 mL of sterile water and 1.5 mL of NaHCO_3_ solution (1M) were then added to the reactor 2 and the mixtures were transferred to TUBE 2 for HPLC purification. *Cis-* and trans-4-[^18^F]fluorogabapentin were isolated by semi-preparative HPLC using a XBridge 5μm C18 prep-column (10 x 250 mm) with a mobile phase of NaH_2_PO_4_ EtOH-H_2_O (10mM, v/v = 5:95, pH = 8) at flow rate of 4 mL/min. The final products were injected on an analytical HPLC (conditions) to assess chemical and radiochemical purity. Since these compounds have no UV absorption in the 210-280 nm range, coinjection with authentic reference standard was not performed. The confirmation of the tracer identity was based on the use of an authentic isomerically pure precursor previously reported and our prior finding that starting from the isomerically pure precursor produces the corresponding isomerically pure product^26^. Likewise, the molar activity could not be calculated from the UV absorption and was estimated based on the molar activity of other ^18^F-labeled compounds produced using the same fluoride source and no-carrier added methods.

## Animal experiments

### Compliance

All experiments involving non-human primates were performed in accordance with the U.S. Department of Agriculture (USDA) Animal Welfare Act and Animal Welfare Regulations (Animal Care Blue Book), Code of Federal Regulations (CFR), Title 9, Chapter 1, Subchapter A, Part 2, Subpart C, §2.31. 2017. Animal experiments were approved by the Animal Care and Use Committee at the Massachusetts General Hospital (MGH).

### Non-human primates (NHPs)

Two male adult rhesus macaques (Monkey 1 and Monkey 2) were used in this study (ages: 13 and 17 years old, respectively). Animal body weights on the days of imaging were 13.26 kg (baseline scan) and 12.78 kg (blocking scan) for Monkey 1 and 15.96 kg (baseline scan) and 14.54 kg (blocking scan) for Monkey 2.

### PET/CT imaging of NHPs

Two rhesus macaques (Monkey 1 and Monkey 2) were scanned on a Discovery MI (GE Healthcare) PET/CT scanner. Prior to PET/CT scan, monkeys were sedated with ketamine/xylazine (10/0.5 mg/kg IM) and were intubated for maintenance anesthesia with isoflurane (1-2% in 100% O_2_). A venous catheter was placed in the saphenous vein for radiotracer injection and an arterial catheter was placed in the posterior tibial artery for blood sampling. The animal was positioned on a heating pad on the bed of the scanner for the duration of the study. A CT scan was acquired prior each PET acquisition for attenuation correction. PET data were acquired in 3D list mode for 180 min following injection of radiotracers. Radiotracers were administered *via* the lateral saphenous vein over a 3-minute infusion and followed by a 3-minute infusion of a saline flush. For the blocking scans, gabapentin (4.7 mg/kg for the [^18^F]tGBP4F scan and 5.0 mg/kg for the [^18^F]cGBP4F scan in 10 mL of saline) was administered 30 min prior to the administration of the radiotracer followed by 10 mL saline flush. All injections were performed using syringe pumps (Medfusion 3500). Dynamic PET data were reconstructed using a validated fully 3D time-of-flight iterative reconstruction algorithm using 3 iterations and 34 subsets while applying corrections for scatter, attenuation, deadtime, random coincident events, and scanner normalization. For all dynamic scans, list mode data were framed into dynamic series of 6×10, 8×15, 6×30, 8×60, 8×120s and remaining were 300 s frames. Final reconstructed images had voxel dimensions of 256 × 256 × 89 and voxel sizes of 1.17 × 1.17 × 2.8 mm^3^.

### Magnetic resonance imaging

For each monkey, a three-dimensional structural T1-weighted magnetization-prepared rapid gradient-echo (MEMPRAGE) magnetic resonance (MR) image was acquired using a 3T Biograph mMR (Siemens Medical Systems) for anatomical reference. Acquisition parameters were as follows: repetition time = 2530 ms; echo time = 1.69 ms; inversion time = 1100 ms; flip angle = 7; voxel size = 1 X 1 X 1 mm^3^; matrix size = 256 X 256 X 176; number of averages = 4.

### Arterial blood sampling and analysis of radiometabolites

Arterial blood sampling was performed during the PET acquisitions. Samples of 1 to 3 mL were initially drawn every 30 seconds upon radiotracer injection and decreased in frequency to every 15 minutes toward the end of the scan. Radiotracer metabolism was characterized from blood samples acquired at 5, 10, 15, 30, 60, 90, 120 and 180 minutes. An additional blood sample of 3 mL was drawn prior tracer injection in order to measure the plasma free fraction (*f*_*p*_). The *f*_p_ was measured by spiking the blood sample with a known amount of radioactivity, gently mixing it and immediately measuring the concentration in measured in whole-blood (WB) and subsequently in plasma (PL). Radioactivity concentration (kBq/cc) was measured in WB and PL following the centrifugation of WB using a calibrated single well gamma counter. Radiometabolite analysis was performed using an automated column switching radioHPLC system^29,30^. Eluent was collected in 1-minute intervals and assayed for radioactivity using a Wallac Wizard gamma counter (1470 model). The procedure adopted for these measurements was similar to the one described previously^31,32^, except for the mobile phases used. Injected plasma samples were initially trapped on the catch column using a mobile phase consisting of either 99:1 10 mM ammonium bicarbonate pH 8 in water:MeCN at 1.8 mL/min and the catch column was backflushed with 95:5 10 mM ammonium phosphate pH 8 in water:EtOH at 1 mL/min and the sample directed onto the analytical column. SUV-radioactivity time courses in WB and in PL were generated by correcting radioactivity concentrations (C [kBq/ml]) for injected dose (ID [MBq]) and animal body weight (BW [kg]) and were calculated as SUV(t) = C(t)/(ID/BW). Measured time courses of percent parent in plasma (%PP(t)) were fitted with a single exponential decay plus a constant. Individual metabolite-corrected arterial input functions (*C*_*p*_ *(t)*) were derived by correcting the total PL radioactivity concentration 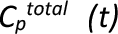 for radiotracer metabolism using individual 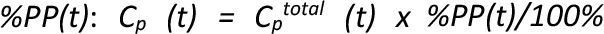^31,32^. The plasma free fractions were measured in triplicate by ultrafiltration following a procedure described previously^31,32^. Briefly, arterial plasma samples of 200 microliters, drawn before radiotracer injection, were spiked with ∼444 kBq of [^18^F]tGBP4F or [^18^F]cGBP4F. Following a 15 min incubation period, radioactive plasma samples were loaded on ultrafiltration tubes (Millipore Centrifree) and centrifuged at 1,500 g for 15 min at room temperature. *f*_*p*_ was calculated as the ratio of free ultrafiltrate to plasma concentration and corrected for binding to the ultrafiltration tube membrane.

### Image registration and processing

All PET processing was performed with an in-house developed Matlab software that uses FSL^33^. Individual brain MR and PET images were aligned into the MRI National Institute of Mental Health macaque template (NMT)^34^ according to the following procedure. An early summed PET image (0-10 min post tracer injection) was rigidly co-registered to the animal’s individual MEMPRAGE MR image which was subsequently registered into the NMT template space using a 12-parameter affine transformation followed by non-linear warping. The transformation matrices were then combined and applied inversely on the rhesus atlases in order to warp all atlases into the native PET image space for extraction of TACs. The composite regions of interest (ROIs) were derived from the two sets of digital anatomical atlases aligned to the NMT space. These were *Paxinos et al*. rhesus brain regions^35^ previously aligned to the *McLaren et al*. rhesus MRI brain template^36^ as well as the INIA19 NeuroMaps rhesus atlas^37^. Briefly, the T1 weighted average MR images of each template were transformed to the NMT template using a 12-parameter affine registration and non-linear warping both weighted on the brain (i.e. after skull stripping). The obtained transformation matrices were then applied to the composite ROIs. Regional TACs were extracted from the native PET image space for the following ROIs: amygdala, caudate, cerebellum grey, insula, nucleus accumbens, occipital gyrus, putamen, parietal lobe, temporal lobe, anterior cingulate, striatum, cerebellum no vermis (whole), cingulate, cortex (all), frontal cortex, hippocampus, occipital lobe, thalamus, white matter and whole brain.

### Brain kinetic analysis

Extracted brain TACs were analyzed by compartmental modeling with reversible one- (1T) and two- (2T) tissue compartment models using the measured metabolite-corrected arterial plasma input function. Each model was assessed with a vascular fraction *V*_*f*_ fixed to 5% and also with *V*_*f*_ estimated as an additional model parameter. For quantification based on compartment models, we followed the consensus nomenclature for in vivo imaging of reversibly binding radioligands^38^. The primary outcome of interest was the total volume of distribution *V*_T_ representing the equilibrium ratio of tracer concentration in tissue relative to its plasma concentration and which is linearly related to the tracer binding to the target. *V*_T_ was calculated as K_1_/k_2_ for a 1T compartment model and as (K_1_/k_2_) x (1 + k_3_/k_4_) for a 2T model. The Logan and multilinear 1 (MA1) graphical analysis methods utilizing arterial input function^39^ were also tested as alternative techniques for estimation of *V*_T_.

### Safety

Experiments involving handling of hazardous chemicals were performed by trained personnel wearing personal protective equipment including lab coats and goggles. Experiments involving radioactive materials were performed by trained personnel wearing personal dosimeters and following ALARA principles. All experiments involving radioactive materials were conducted according to an approved radioactive materials permit. Experiments involving handling of animal fluids were performed by trained personnel wearing appropriate personal protective equipment and following institutional approved biosafety protocols.

## ABBREVIATIONS

1TCM: 1 tissue-compartment model
cGBP4F: *cis*-4-fluorogabapentin
GBP: gabapentin
MA1: multilinear analysis-1
MRI: magnetic resonance imaging
PET: positron emission tomography
ROI: region of interest
TAC: time-activity curves
tGBP4F: *trans*-4-fluorogabapentin

## ACKNOWLEDGEMENTS

We thank David Lee and Timothy Beaudoin at the MGH Gordon PET cyclotron facility for producing fluorine-18. We thank the veterinary staff (Helen Deng) for assistance with animal handling. Funding: This study was supported by R21NS120139 (PB), K99EB033407 (YPZ), S10OD018035 (GEF, MDN), P41EB022544 (GEF, MDN) and MGH Fund for Medical Discovery (YPZ).

## AUTHOR CONTRIBUTIONS

YPZ contributed to the study design, developed the automated radiosynthesis protocols, and synthesized the radiotracers; MDN: contributed to the study design, performed the PET imaging in monkeys and contributed to data interpretation; VB assisted with the PET imaging and performed initial data analysis; MTMS, YS, SHM processed and analyzed the blood samples; GEF: contributed to data interpretation; NJG processed and analyzed the PET data assisted by MTMS; PB contributed to the study design, radiosynthesis method development and data analysis and interpretation. YPZ, VB, NJG and PB wrote the manuscript and all authors reviewed and approved it.

## SUPPLEMENTARY MATERIAL AND DATA AVAILABILITY

Supplemental material for this article including 7 supplemental figures is available online. The datasets generated and/or analyzed during the current study are available from the corresponding authors on reasonable request.

