## Supplementary material for "Evaluation of *trans*- and *cis*-4-[^18^F]fluorogabapentin for brain PET imaging": Sup. Fig.

| Item | Description | Page |
|---|---|---|
| Figure S1 | Synthesizer flow diagram and reagents | S3 |
| Figure S2 | Analytical chromatograms (UV and radio) of [$^{18}$F]cGBP4F and [$^{18}$F]tGBP4F | S4 |
| Figure S3 | Biexponential fits of radioactivity concentration in blood | S5 |
| Figure S4 | Whole blood to plasma ratio | S6 |
| Figure S5 | Radiometabolites in blood | S7 |
| Figure S6 | Logan graphical analysis fits | S8 |
| Figure S7 | Correlation $V_T$ calculated via Logan and MA-1 with $V_T$ from 1TCM. | S9 |

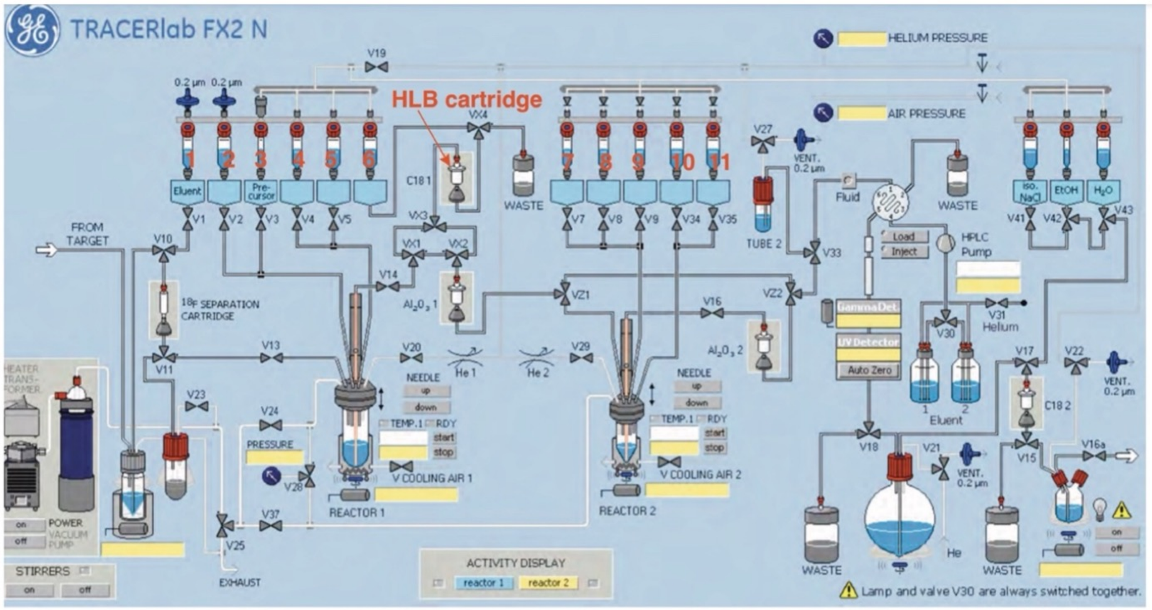


| Vials | Reagent | Vials | Reagent |
|---|---|---|---|
| 1 | 1mL of acetonitrile/water (1:1) solution of Kryptofix 2.2.2 ($K_{222}$, 15 mg) and $K_2CO_3$ (1.25 mg) | 7 | 0.3 mL of NaOH solution (5N) |
| 2 | 6 mL of sterile water | 8 | 0.6 mL of HCl solution (6N) |
| 3 | 0.3 mL of anhydrous DMSO solution of precursor | 9 | 0.2 mL of NaOH solution (5N) |
| 4 | 1 mL of anhydrous acetonitrile | 10 | 2 mL of sterile water |
| 5 | 6 mL of sterile water | 11 | 1.5 mL of $NaHCO_3$ solution (1M) |
| 6 | 2 mL of acetonitrile | | |

**Figure S1. Synthesizer flow diagram and reagents**. Flow diagram of the synthesis module used in the synthesis (GE TRACERlab FX2 N) and table of reagents.

A

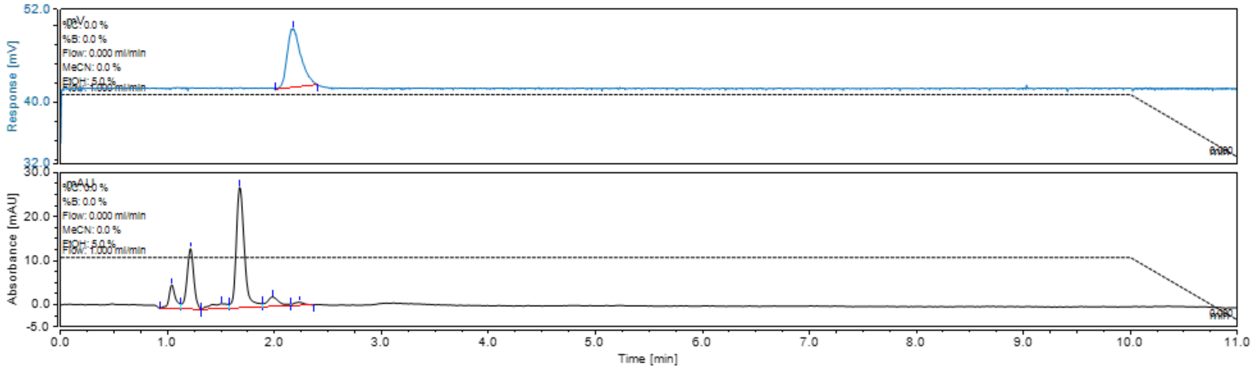


B

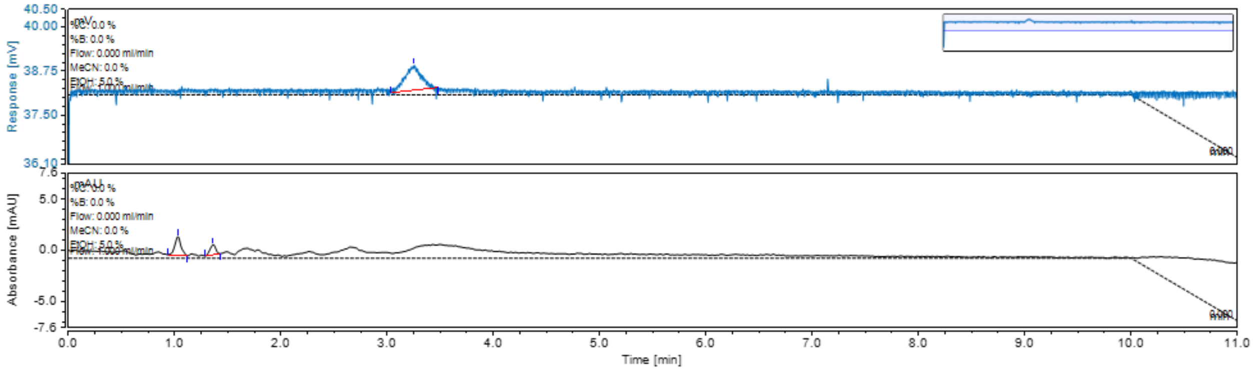


**Figure S2.** Analytical chromatograms (top: radio-detector; bottom UV detector at 210 nm) of [$^{18}$F]cGBP4F (**A**) and [$^{18}$F]tGBP4F (**B**).

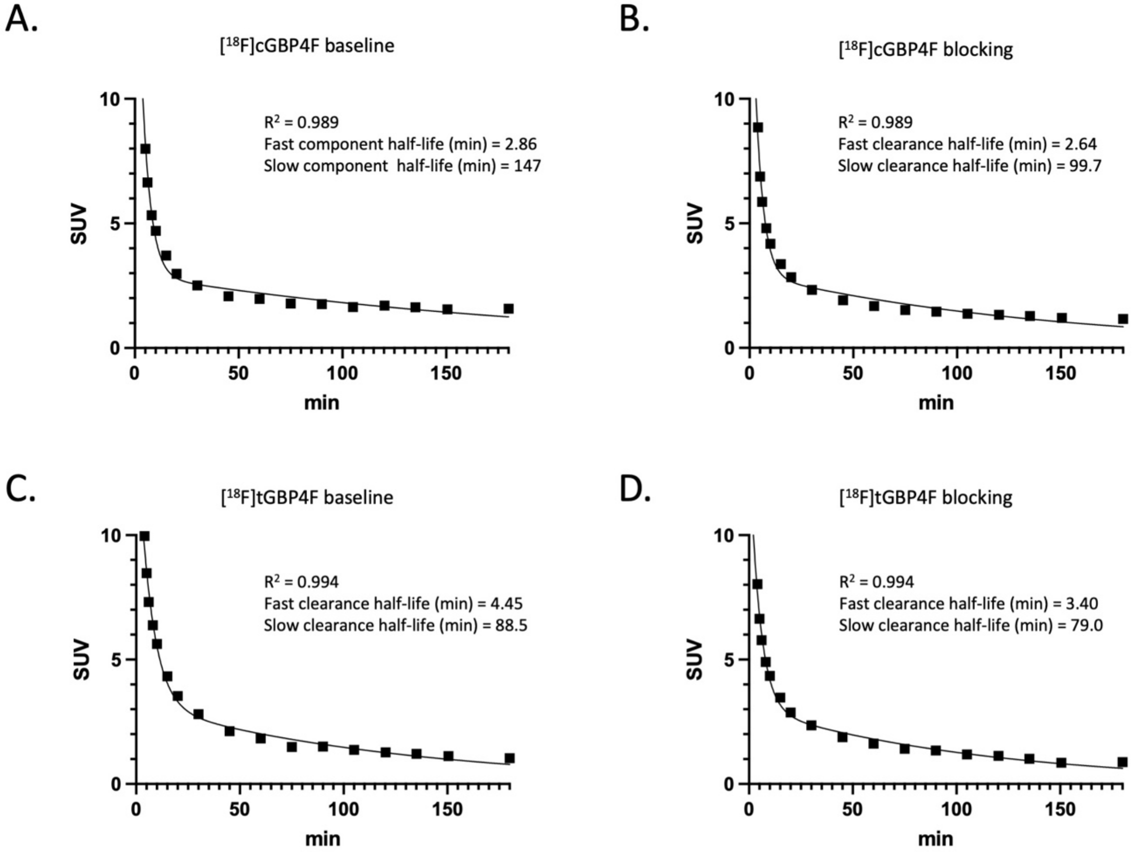


**Figure S3**. Biexponential fits of plasma parent concentration in blood. **A.** [$^{18}$F]cGBP4F baseline. **B.** [$^{18}$F]cGBP4F blocking. **C.** [$^{18}$F]tGBP4F baseline. **D.** [$^{18}$F]tGBP4F blocking.

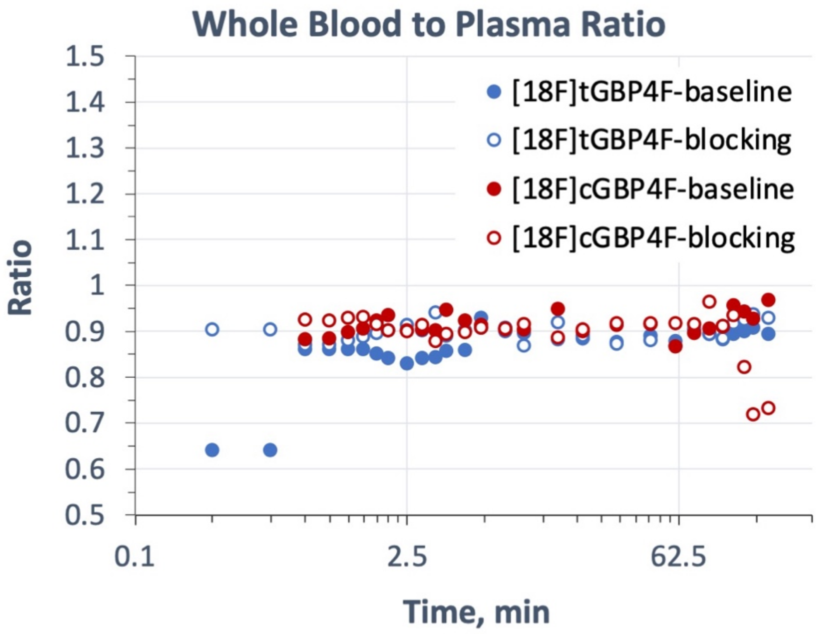


**Figure S4.** Whole blood to plasma ratio

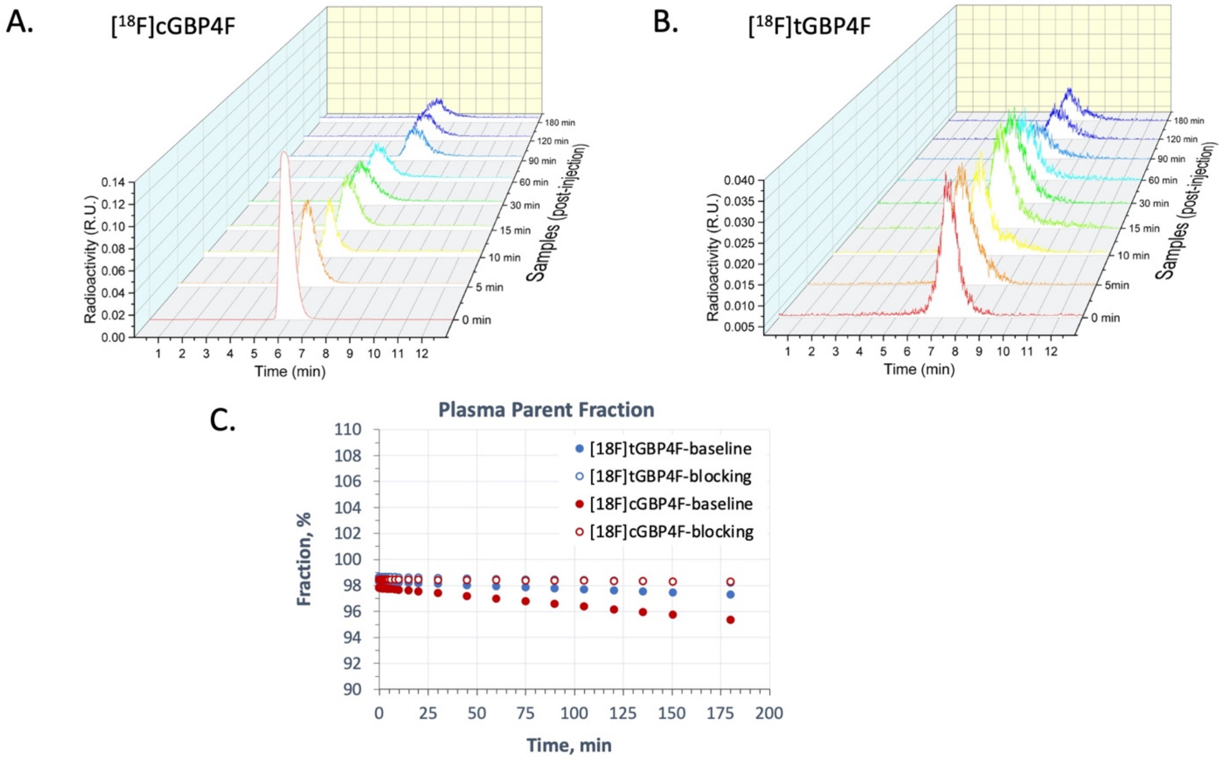


**Figure S5. Radiometabolites in blood.** Representative HPLC radiochromatograms of plasma samples of [$^{18}$F]cGBP4F (**A**) and [$^{18}$F]tGBP4F (**B**) obtained at different times post injection (R.U.: relative units). **C**. Percent parent fraction in plasma.

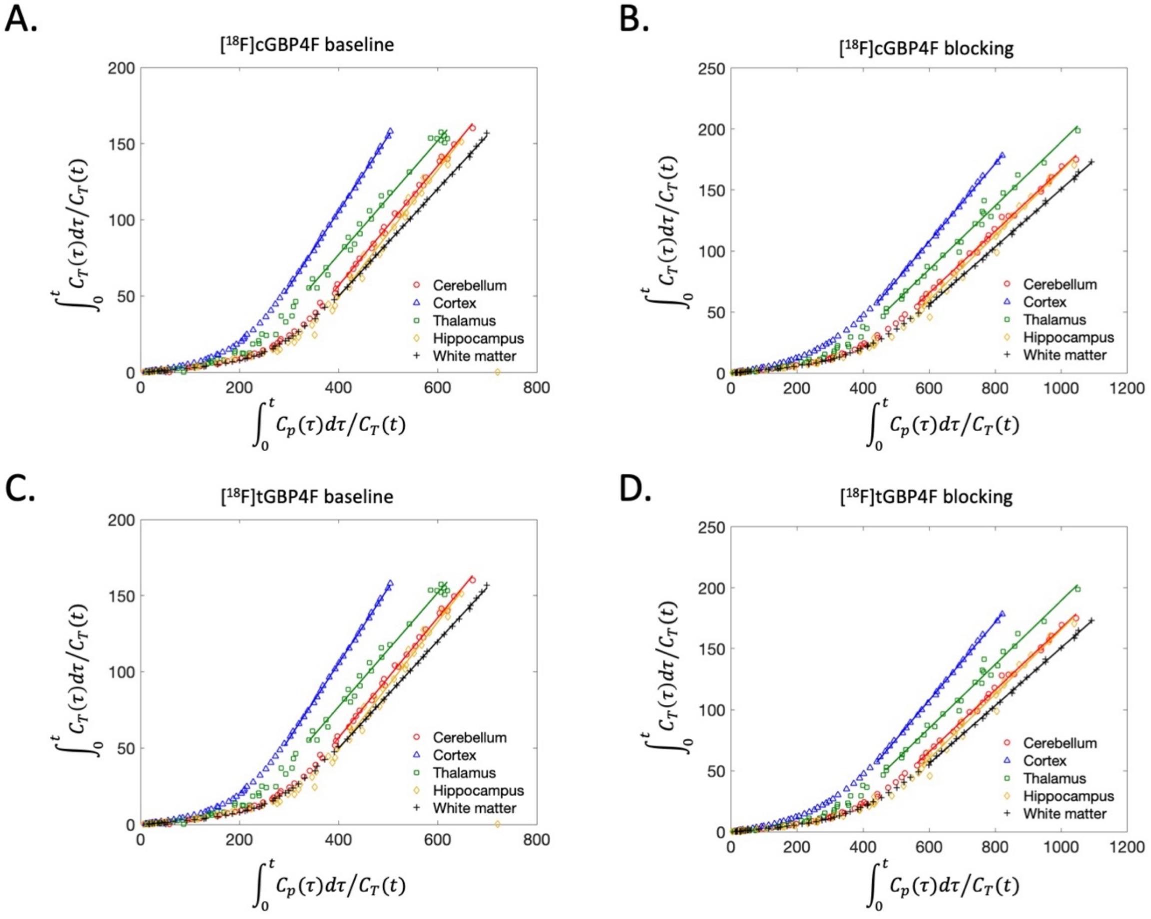


**Figure S6**. Logan graphical analysis. **A.** [$^{18}$F]cGBP4F baseline. **B.** [$^{18}$F]cGBP4F blocking. **C.** [$^{18}$F]tGBP4F baseline. **D.** [$^{18}$F]tGBP4F blocking.

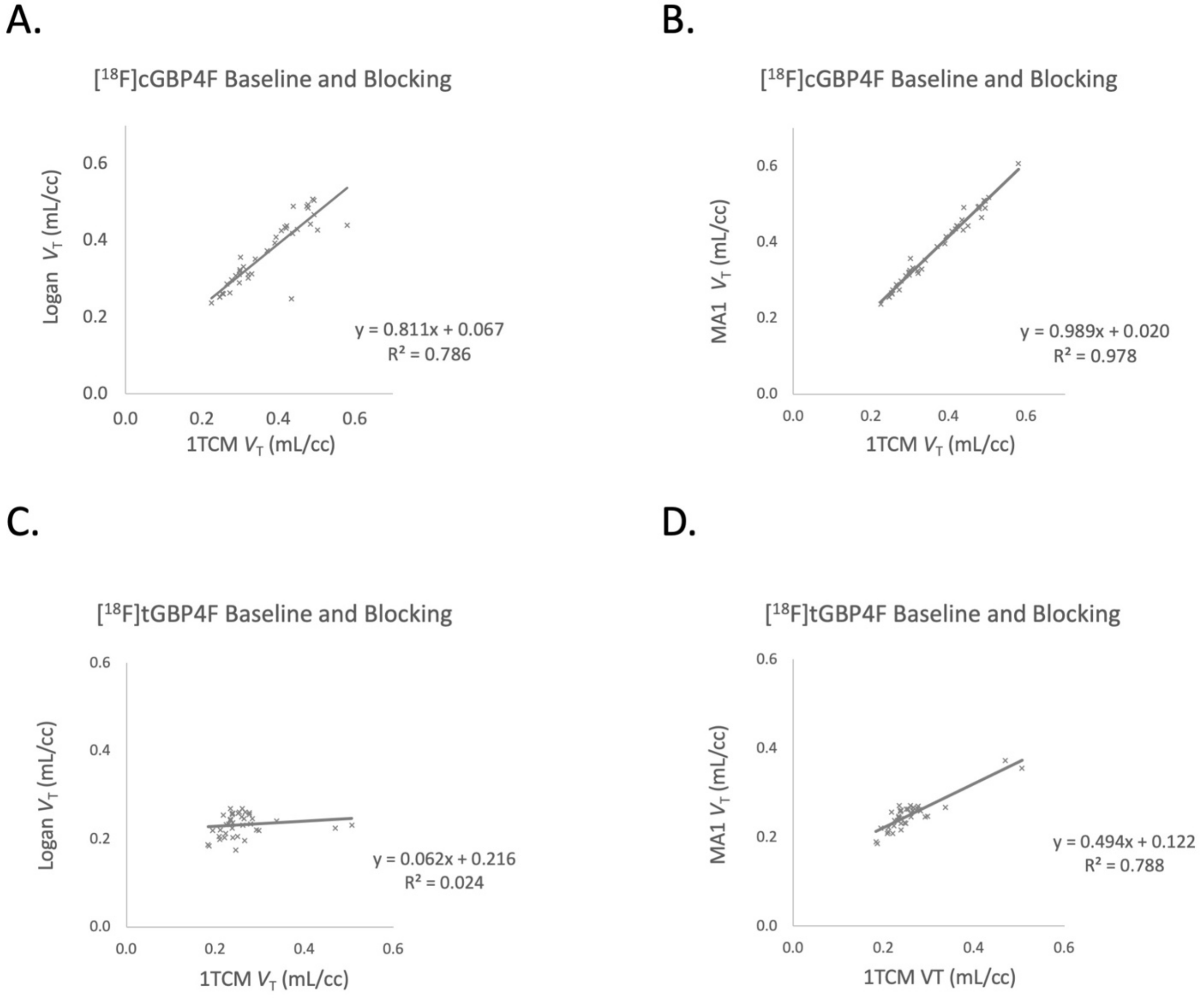


**Figure S7**. Correlation $V_T$ calculated via Logan and MA-1 with $V_T$ from 1TCM. **A.** [$^{18}$F]cGBP4F baseline and blocking Logan – 1TCM. **B.** [$^{18}$F]cGBP4F baseline and blocking MA1 – 1TCM. **C.** [$^{18}$F]tGBP4F baseline and blocking Logan – 1TCM. **D.** [$^{18}$F]tGBP4F baseline and blocking MA1 – 1TCM.
